# A naturally occurring frameshift mutation in the UNUSUAL FLORAL ORGANS gene associated with the marimo floral phenotype in gerbera

**DOI:** 10.64898/2026.08.09.743735

**Authors:** Taro Hattori, Riki Shimada, Moe Nagakura, Ryo Ando, Sachiko Isobe, Naoyuki Tajima, Hideki Hirakawa, Kenta Shirasawa, Akiyoshi Tominaga

## Abstract

**Background:** The capitulum of Asteraceae is a highly specialized inflorescence whose formation requires the coordinated regulation of multiple developmental processes, including floral organ identity and floral meristem determinacy. The LEAFY (LFY)–UNUSUAL FLORAL ORGANS (UFO) regulatory module is known to play an important role in flower development; however, naturally occurring mutations affecting this pathway have not been genetically characterized in gerbera (*Gerbera hybrida*).

**Results:** In this study, we characterized a novel gerbera mutant identified during a commercial crossing program and named it *marimo* based on its green, spherical capitulum. Morphological observations revealed the repeated formation of secondary and tertiary floret-like organs within primary floret-like organs. Scanning electron microscopy showed that the epidermal structure of the green organs in *marimo* was similar to that of wild-type involucral bracts. RNA sequencing identified numerous differentially expressed genes between marimo and the wild type, and network and Gene Ontology analyses highlighted gene groups associated with flower development, reproductive organ differentiation, and tissue structure formation. RNA-seq analysis showed increased expression of LFY and reduced expression of GGLO1, a PISTILLATA/GLOBOSA-like B-class MADS-box gene, in the marimo mutant. RT-qPCR analysis of a segregating population further confirmed reduced GGLO1 expression in marimo-type individuals. In addition, a single-nucleotide deletion was identified in the coding region of UFO. This deletion was predicted to cause a frameshift and a premature stop codon. In selfed progeny of No. 251, the UFO genotype was fully associated with capitulum phenotype, and only individuals homozygous for the mutant allele exhibited the *marimo* phenotype.

**Conclusions:** These results indicate that the naturally occurring frameshift mutation in UFO is the strongest candidate variant underlying the *marimo* phenotype. RNA-seq analysis showed increased LFY expression and markedly reduced GGLO1 expression in the marimo mutant. Reduced activity of the LFY–UFO regulatory module may therefore have altered the expression of GGLO1 and other floral organ development-related genes despite the continued expression of LFY. These changes may have affected both floral organ identity and floral meristem determinacy, resulting in the formation of green involucral bract-like organs and the repeated production of floret-like organs. The *marimo* mutant provides a useful genetic resource for investigating capitulum development in Asteraceae and may also serve as breeding material for introducing novel ornamental traits into gerbera.

## Introduction

Gerbera (*Gerbera hybrida*) is one of the world’s most important ornamental crops and has considerable horticultural value because of its extensive diversity in flower color, floral morphology, and capitulum architecture (Kloos et al., 2004; Aoyagi et al., 2026). Numerous cultivars with diverse floral forms have been developed through breeding, and gerbera is widely used not only as breeding material for ornamental plants but also as a model system for investigating the mechanisms of flower development (Yu et al., 1999; Broholm et al., 2010). Because floral morphology is a major determinant of ornamental value, understanding the molecular mechanisms that regulate capitulum development is important for both fundamental plant science and horticultural breeding. More recently, chromosome-level genome sequences have become available for the Asteraceae model species *Chrysanthemum seticuspe* (Nakano et al., 2021) and gerbera (Aoyagi et al., 2026), providing a foundation for the molecular analysis of complex traits in Asteraceae.

Gerbera belongs to Asteraceae, one of the largest families of flowering plants, which comprises approximately 10% of all angiosperm species (Funk et al., 2009; Zhang and Elomaa, 2024). A defining morphological feature of Asteraceae is the capitulum. This highly specialized inflorescence consists of numerous densely arranged florets and appears as a single flower, forming a pseudanthium that is considered one of the major morphological innovations underlying the evolutionary success of Asteraceae (Broholm et al., 2014; Elomaa et al., 2018; Gurung et al., 2024; Zhang and Elomaa, 2024). The gerbera capitulum consists of ray florets at the periphery, trans florets in the intermediate region, and disc florets at the center, arranged in concentric patterns (Yu et al., 1999; Broholm et al., 2014; Zhang et al., 2021). The formation of this complex reproductive structure requires the coordinated regulation of inflorescence meristem identity, floral meristem identity, floral organ identity, and floral meristem determinacy (Elomaa et al., 2018; Gurung et al., 2024).

Flower development in angiosperms is controlled by highly conserved gene regulatory networks. In the ABC model and its extended forms, floral organ identities, including those of sepals, petals, stamens, and carpels, are specified by combinations of floral organ identity genes (Coen and Meyerowitz, 1991; Bowman et al., 1991; Pelaz et al., 2000; Ditta et al., 2004). In addition to these floral organ identity genes, floral meristem identity genes initiate flower formation and regulate subsequent developmental programs (Weigel et al., 1992; Weigel and Meyerowitz, 1994). Normal flower development also requires floral meristem determinacy, which terminates meristem activity after a defined number of floral organs have formed (Ratcliffe et al., 1999). Disruption of these regulatory processes can result in homeotic transformation of floral organs and reiterative organ formation (Bowman et al., 1991; Ratcliffe et al., 1999).

Among the key regulators of flower development, LEAFY (LFY) is a central transcription factor controlling floral meristem identity. LFY promotes the transition from inflorescence meristem to floral meristem and activates numerous downstream genes involved in floral organ formation (Weigel et al., 1992; Weigel and Meyerowitz, 1994). UNUSUAL FLORAL ORGANS (UFO), which encodes an F-box protein, functions together with LFY in *Arabidopsis* to activate the B-class floral organ identity genes APETALA3 (AP3) and PISTILLATA (PI), thereby regulating petal and stamen development (Levin and Meyerowitz, 1995; Wilkinson and Haughn, 1995; Lee et al., 1997; Hepworth et al., 2006; Chae et al., 2008). Thus, LFY expression alone may not be sufficient for the full activation of some B-class genes, and functional cooperation between LFY and UFO is considered necessary. The LFY–UFO regulatory module therefore represents a central pathway linking floral meristem identity with floral organ development.

In gerbera, the LFY–UFO regulatory module has also been recruited for capitulum development. RNA interference-mediated suppression of either GhLFY or GhUFO causes severe abnormalities in capitulum structure, including the replacement of normal floral organs by green involucral bract-like organs and the repeated formation of secondary and tertiary floret-like organs (Zhao et al., 2016). These findings indicate that GhLFY and GhUFO contribute not only to floral organ identity but also to floral meristem identity and determinacy.

The PISTILLATA/GLOBOSA-like B-class MADS-box gene GGLO1 also plays an important role in petal and stamen identity in gerbera. Suppression of GGLO1 expression causes homeotic changes in petals and stamens, indicating that GGLO1 is a major component of the B-class floral organ identity program in gerbera (Broholm et al., 2010). Although direct regulation of GGLO1 by the LFY–UFO module has not been demonstrated in gerbera, the ability of LFY and UFO to activate PI in *Arabidopsis* suggests that GGLO1 may be one of the downstream candidates regulated by this module.

SEPALLATA-like MADS-box genes also play important roles in patterning the gerbera capitulum (Uimari et al., 2004; Zhang et al., 2017). These findings suggest that conserved regulators of flower development have been evolutionarily recruited to control the specialized architecture of the Asteraceae capitulum (Elomaa et al., 2018; Zhang and Elomaa, 2024).

Most of these findings, however, have been obtained from transgenic plants in which target gene expression was experimentally suppressed. Naturally occurring gerbera mutants in which the LFY–UFO regulatory module is disrupted have not yet been subjected to detailed genetic analysis.

Against this background, a gerbera individual with a distinctive green, spherical capitulum was recently identified among progeny derived from a cross between No. 251 and an unknown breeding line during a commercial breeding program at Green Tech Co., Ltd. This individual, which was analyzed in the present study, was named *marimo* because its capitulum consisted of numerous densely packed green organs and resembled the spherical green alga marimo (*Aegagropila linnaei*) found in Japanese lakes. The capitulum of this mutant contained repeated secondary and tertiary floret-like organs within primary floret-like organs, suggesting defects in both floral organ identity and floral meristem determinacy. Individuals with similar phenotypes were subsequently observed among selfed progeny of No. 251. In addition, similar phenotypes segregated in progeny from reciprocal crosses between No. 251 and No. 3065.

In this study, we investigated the morphological and molecular genetic basis of the *marimo* phenotype through detailed morphological characterization, scanning electron microscopy of epidermal structures, RNA-seq-based transcriptome analysis, gene network analysis, and nucleotide sequence analysis of candidate gene. We further examined the association between candidate genotypes and capitulum phenotypes in selfed progeny of No. 251 and No. 3065 and evaluated phenotypic segregation in reciprocal crosses between No. 251 and No. 3065.

## Materials and methods

### Plant materials

The *marimo* mutant used in this study was identified during a commercial breeding program for cut-flower gerbera at Green Tech Co., Ltd. The original *marimo* mutant was obtained from progeny derived from a cross between No. 251 and an unknown breeding line. Individuals exhibiting a similar *marimo* phenotype were also identified among selfed progeny of No. 251 and progeny from reciprocal crosses between No. 251 and No. 3065.

The plant materials used in this study included the *marimo* mutant derived from the cross between No. 251 and the unknown breeding line, No. 251, No. 3065, and the wild-type cultivar ‘Opal’, for which a whole-genome reference sequence is available (Aoyagi et al., 2026) (Fig. 1).

**Figure 1.**
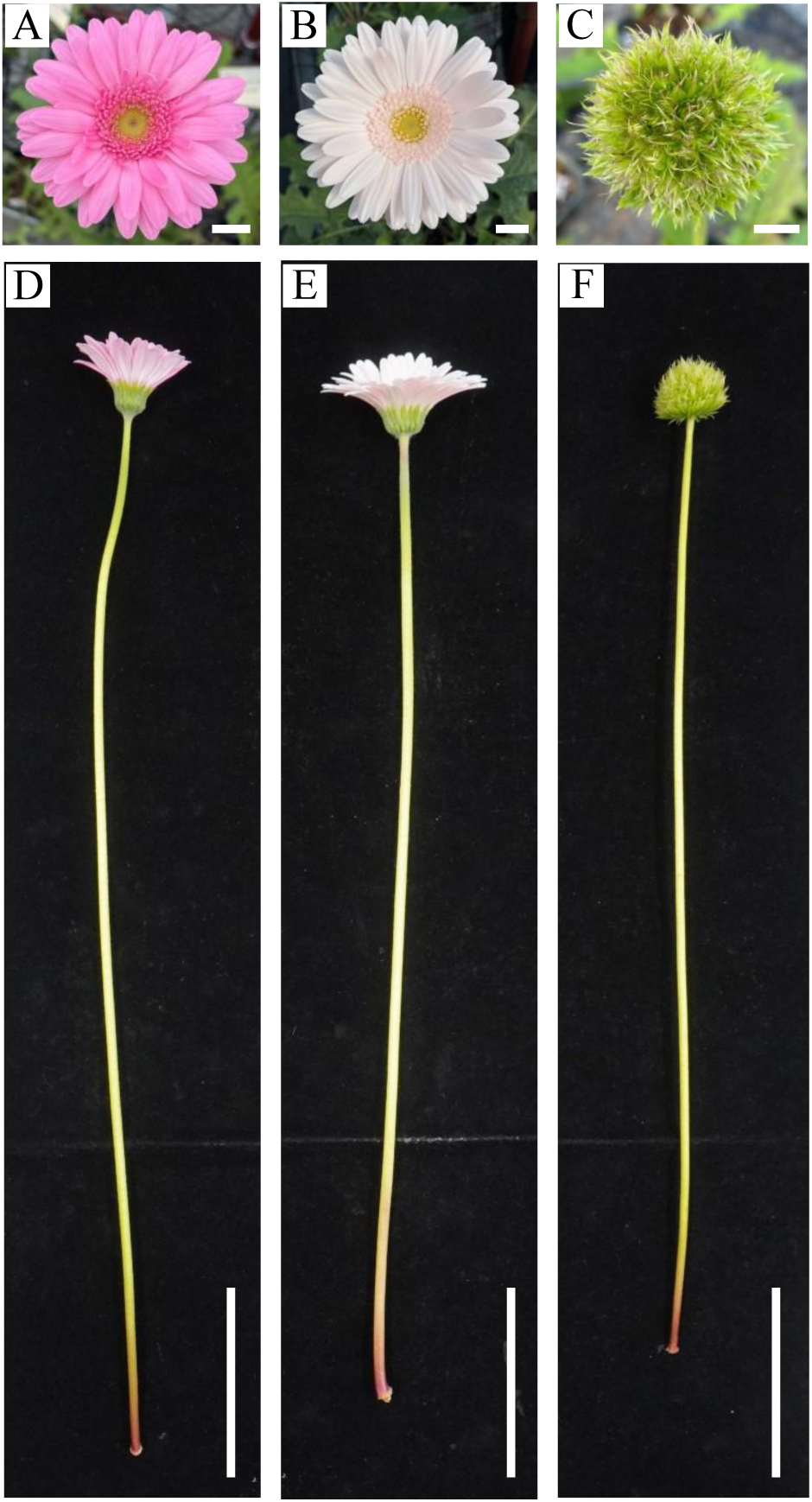
Morphological comparison of wild-type gerbera and the *marimo* mutant. (A–C) Capitula of the wild-type line No. 251 (A), the wild-type cultivar ‘Opal’ (B), and the *marimo* mutant (C). (D–F) Whole cut flowers of No. 251 (D), ‘Opal’ (E), and the *marimo* mutant (F). Scale bars: A–C, 1 cm; D–F, 10 cm.

For morphological observations and molecular analyses, 15 two-year-old *marimo* plants, eight two-year-old No. 251 plants, and 15 two-year-old ‘Opal’ plants were used. Each plant was grown in a No. 8 pot filled with a 1:1 mixture of commercial potting soil and vermiculite. Fertilization and removal of senescent or excessive leaves were performed once every two weeks. The plants were cultivated in a greenhouse at the Fujieda Field, Center for Education and Research in Field Sciences, Faculty of Agriculture, Shizuoka University. During winter, the greenhouse was heated to maintain the temperature above 15°C.

For genetic segregation analysis, 40 selfed progeny of No. 251 and eight selfed progeny of No. 3065 were examined. The occurrence of the *marimo* phenotype was also investigated in 34 progeny from a cross using No. 3065 as the seed parent and No. 251 as the pollen parent, and in 28 progeny from the reciprocal cross using No. 251 as the seed parent and No. 3065 as the pollen parent.

### Crossing and phenotypic evaluation of progeny

Self-pollination and reciprocal crosses were performed using No. 251 and No. 3065. The reciprocal crosses consisted of No. 3065 as the seed parent and No. 251 as the pollen parent, and No. 251 as the seed parent and No. 3065 as the pollen parent. Capitula at anthesis were used for crossing, and pollen collected from the pollen parent was manually applied to the stigmas of the seed parent. The resulting seeds were sown and the plants were grown until flowering. Progeny were classified as wild type or *marimo* type based on capitulum morphology.

### Morphological observation

Green organs constituting the capitula of the *marimo* mutant, as well as involucral bracts, leaves, and pappi of ‘Opal’ and No. 251, were collected. Their morphology was compared using a stereomicroscope (SZX7, Olympus, Tokyo, Japan), and fresh samples were mounted directly on aluminum stubs without fixation, dehydration, or metal coating and were observed under low-vacuum conditions using a scanning electron microscope (Miniscope TM3030Plus, Hitachi High-Tech, Tokyo, Japan). The adaxial and abaxial surfaces of involucral bracts and leaves were examined. In addition, capitula of the *marimo* mutant were dissected, and the formation patterns of primary, secondary, and tertiary floret-like organs were examined in detail under a stereomicroscope.

### RNA extraction and RNA sequencing

For RNA-seq analysis, unopened developing capitula approximately 15 mm in diameter were collected from the *marimo* mutant, No. 251, and ‘Opal’. Three unopened developing capitula were collected from three independent plants of each genotype, with one capitulum from each plant used as a biological replicate. Samples were immediately frozen in liquid nitrogen and stored at −80°C until RNA extraction. The frozen tissues were pulverized using a ShakeMan homogenizer, and total RNA was extracted using the RNeasy Plant Mini Kit (QIAGEN, Hilden, Germany). The extracted RNA was treated with RNase-free DNase I (New England Biolabs, Ipswich, MA, USA). RNA-seq libraries were prepared using the TruSeq Stranded mRNA HT Sample Prep Kit (Illumina). RNA sequencing was outsourced to the Kazusa DNA Research Institute and performed on a DNBSEQ-G400RS platform to generate 100 bp paired-end reads.

### RNA-seq data analysis

The quality of the raw reads was assessed using FastQC (Andrews, 2010), and adapter sequences and low quality reads were removed using Trimmomatic v0.39 (Bolger et al., 2014). The quality-filtered reads were mapped to the gerbera reference genome (Aoyagi, Shimada et al., 2026) using HISAT2 v2.2.1 (Kim et al., 2019). Based on the gene model annotations, the number of fragments assigned to each gene was quantified using featureCounts v2.1.1 (Liao et al., 2014).

The resulting count data were normalized and analyzed for differential expression using DESeq2 v1.50.2 (Love et al., 2014). The marimo mutant was compared separately with No. 251 and ‘Opal’. Genes with an absolute log2 fold change of at least 1 (|log2FC| ≥ 1) and a Benjamini–Hochberg-adjusted p-value ≤ 0.05 were considered differentially expressed genes (DEGs). Variance stabilizing transformation (VST) was applied to the count data, and principal component analysis (PCA) was performed using the transformed expression data. The PCA results were visualized using the ggplot2 package in R v4.5.2 (Wickham, 2016; R Core Team, 2025). In addition, the overlap between the DEGs identified in the two comparisons was visualized using a Venn diagram.

### Network analysis and Gene Ontology analysis

Differentially expressed genes commonly identified in the comparisons of the marimo mutant with No. 251 and with ‘Opal’ were used for protein–protein interaction (PPI) network analysis. To identify Arabidopsis thaliana orthologs corresponding to the gerbera genes, bidirectional sequence similarity searches were performed between the predicted protein sequences of the gerbera reference genome and the TAIR10 protein sequences using DIAMOND v2.1.21 (Buchfink et al., 2021). The searches were conducted in more-sensitive mode, with a maximum E-value of 1 × 10⁻³ and minimum alignment coverages of 30% for both the query and subject sequences. Pairs of sequences that were mutually identified as the top hits in the bidirectional searches were extracted as reciprocal best hits (RBHs). The A. thaliana orthologs identified by RBH analysis were submitted to the STRING database (Szklarczyk et al., 2025) to retrieve known and predicted protein–protein interaction information. The resulting network was imported into Cytoscape v3.10.4 (Shannon et al., 2003) for visualization and network analysis. Node colors were determined based on the mean log2 fold change across the two comparisons: the marimo mutant versus No. 251 and the marimo mutant versus ‘Opal’.

Markov clustering (MCL) analysis was subsequently performed using clusterMaker2 v2.3.4 (Utriainen and Morris, 2023) in Cytoscape to partition the PPI network into densely interconnected clusters. The network was treated as an undirected graph, and the STRING combined score was used as the edge weight. The inflation parameter for MCL analysis was set to 4.0. Gene Ontology (GO) enrichment analysis was performed separately for genes in MCL clusters 1–3 using clusterProfiler v4.18.4 (Wu et al., 2021) in R. The top 10 significantly enriched GO terms with both p-values and adjusted p-values ≤ 0.05 and q-values ≤ 0.20 were visualized using the dotplot function in the enrichplot package.

### Quantitative reverse transcription PCR analysis of GGLO1

A segregating population derived from self-pollination of No. 251 was used to analyze GGLO1 expression by RT-qPCR. Total RNA was extracted from developing capitula of each individual using the RNeasy Plant Mini Kit (QIAGEN, Hilden, Germany), and cDNA was synthesized using the PrimeScript RT Reagent Kit (Takara Bio, Shiga, Japan). Quantitative PCR was performed using TB Green Premix Ex Taq II and a Thermal Cycler Dice Real Time System II (Takara Bio). The amplification program consisted of an initial denaturation at 95°C for 30 s, followed by 40 cycles of 95°C for 5 s and 60°C for 30 s. A melting curve analysis was subsequently performed at 95°C for 15 s, 60°C for 30 s, and 95°C for 15 s to confirm the specificity of the amplification products. GGLO1 expression levels were normalized using 2PS as the reference gene, using previously reported primers (Naing et al., 2018). The primer sequences for GGLO1 are listed in Table 1.

**Table 1.**
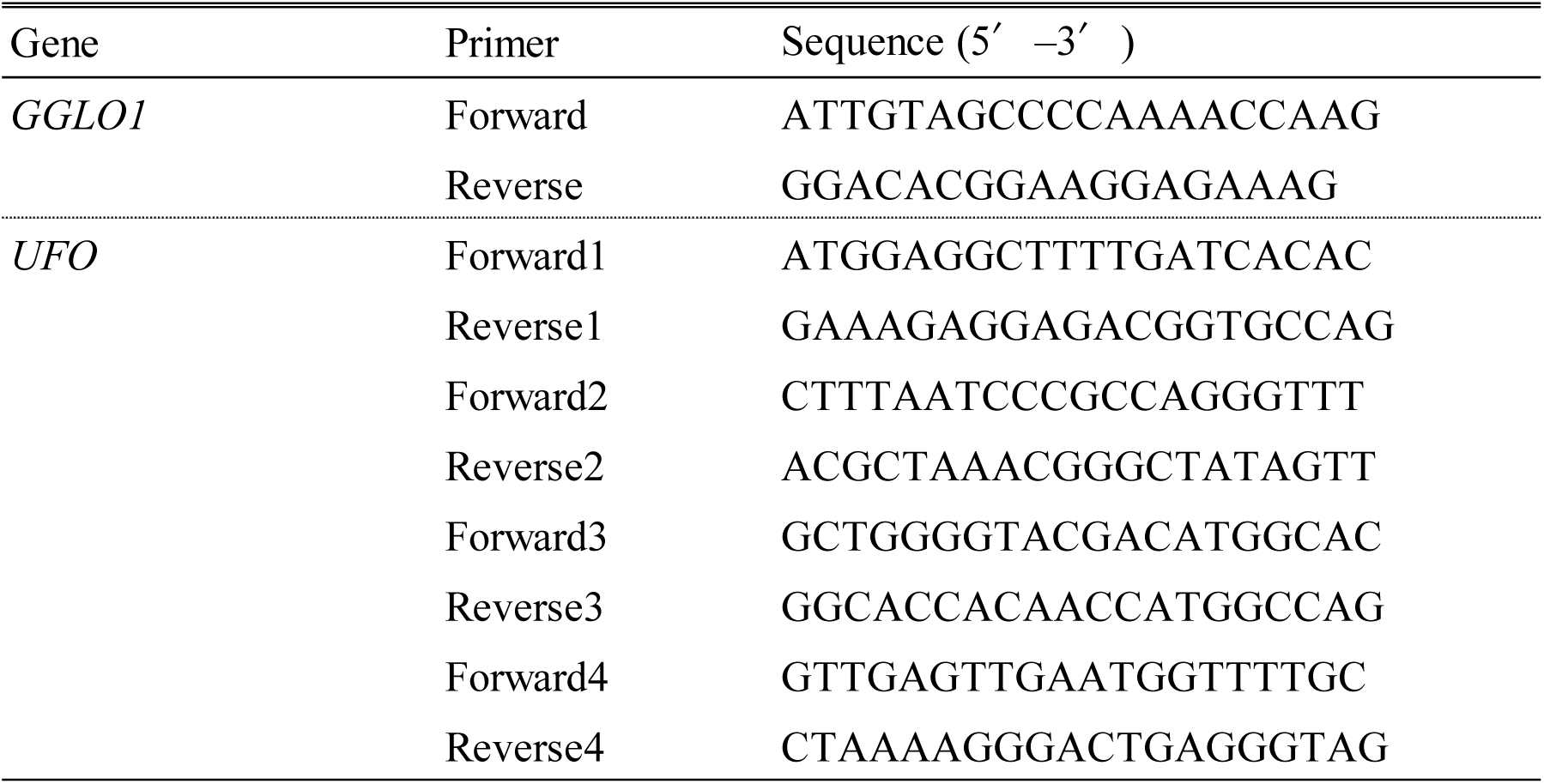
Primer sequences used for GGLO1 expression analysis and UFO sequence and genotype analyses. Forward and reverse primer sequences used for RT-qPCR analysis of GGLO1 and nucleotide sequence and genotype analyses of UFO. Primers were designed based on the sequences of GGLO1 (Ghyb013_g01155.1) and UFO (Ghyb006_g00062.1) in the gerbera reference genome Ghyb_p1.1. Primer sequences are shown in the 5′–3′ direction.

### Nucleotide sequence analysis of UFO

Based on the gene annotation of the gerbera reference genome Ghyb_p1.1 (Aoyagi et al., 2026), the gene model corresponding to UFO (Ghyb006_g00062.1) was selected for analysis. Genomic DNA was extracted from leaves of ‘Opal’, No. 251, No. 3065, and the *marimo* mutant using a cetyltrimethylammonium bromide (CTAB)-based method. Primers were designed based on the sequence of the selected gene model, and the coding region of UFO was amplified by PCR from each plant material. Each 10-µL PCR mixture contained 2× PCR Buffer for KOD FX Neo, 2 mM dNTPs, 10 µM forward primer, 10 µM reverse primer, KOD FX Neo, and 1 µL genomic DNA. PCR amplification was performed using a Thermal Cycler Dice Touch (Takara Bio) for 35 cycles of 95°C for 30 s, 65°C for 30 s, and 72°C for 20 s.

PCR products showing a clear single band after electrophoresis were purified using the NucleoSpin Gel and PCR Clean-up Kit (Macherey-Nagel, Düren, Germany) and subjected to Sanger sequencing. The resulting nucleotide sequences were compared among ‘Opal’, No. 251, No. 3065, and the *marimo* mutant to identify sequence variants.

The UFO coding sequence was also compared with the previously reported gerbera GhUFO sequence (GenBank accession no. KU554695). Deduced amino acid sequences were generated from the obtained UFO nucleotide sequences, and the effects of the single-nucleotide deletion-induced frameshift on protein length and amino acid sequence were predicted. The primer sequences used are listed in Table 1.

### Genetic segregation analysis

To evaluate the association between the single-nucleotide deletion identified in UFO and the marimo phenotype, UFO genotypes and capitulum phenotypes were examined in 40 selfed progeny of No. 251. To further assess the inheritance of the mutant allele, the same analyses were performed using eight selfed progeny of No. 3065.

The UFO genotype of each individual was classified as wild-type homozygous (WT/WT), heterozygous (WT/UFOΔC), or mutant homozygous (UFOΔC/UFOΔC). Capitulum phenotypes were classified as wild type, characterized by the formation of normal ray, trans, and disc florets, or *marimo* type, characterized by a spherical capitulum composed of numerous green organs. The genomic region surrounding the single-nucleotide deletion in UFO was amplified by PCR from each progeny, and the PCR products were subjected to direct Sanger sequencing. UFO genotypes were determined from the sequence chromatograms and classified as wild-type homozygous (WT/WT), heterozygous (WT/UFOΔC), or mutant homozygous (UFOΔC/UFOΔC).

### Statistical analysis

Chi-square goodness-of-fit tests were used to evaluate whether the observed segregation of capitulum phenotypes conformed to the expected 3:1 ratio and whether the segregation of UFO genotypes conformed to the expected 1:2:1 ratio. Statistical analyses were performed using R version 4.5.2, and differences were considered statistically significant at P < 0.05.

## Results

### Occurrence of the *marimo* phenotype during the crossing program

An examination of breeding records from Green Tech Co., Ltd. showed that the original *marimo* mutant was identified among progeny derived from a cross between No. 251 and an unknown breeding line. Individuals exhibiting a similar *marimo* phenotype were subsequently identified among selfed progeny of No. 251 and among progeny from reciprocal crosses between No. 251 and No. 3065. Among 34 progeny from the cross in which No. 3065 was used as the seed parent and No. 251 as the pollen parent, eight exhibited the *marimo* phenotype and 26 exhibited the wild-type phenotype. Among 28 progeny from the reciprocal cross, in which No. 251 was used as the seed parent and No. 3065 as the pollen parent, seven exhibited the *marimo* phenotype and 21 exhibited the wild-type phenotype. In both crossing directions, the observed segregation was consistent with the expected 3:1 ratio of wild-type to marimo-type individuals. The segregation did not differ significantly from the expected ratio in either the No. 3065 × No. 251 cross (χ² = 0.039, P = 0.843) or the reciprocal No. 251 × No. 3065 cross (χ² = 0.000, P = 1.000). These results suggested that both No. 251 and No. 3065 carried a recessive genetic factor associated with the marimo phenotype in the heterozygous state.

### Morphological characteristics of the *marimo* mutant

The *marimo* mutant formed green, spherical, and compact capitula that differed markedly from those of the wild-type line No. 251 and the wild-type cultivar ‘Opal’ (Fig. 1A–C). No. 251 and ‘Opal’ formed capitula consisting of normal ray, trans, and disc florets. In contrast, most of the *marimo* capitulum consisted of elongated green organs, and clearly differentiated petals, stamens, and pistils were not observed. The *marimo* mutant produced elongated scapes similar to those of the wild types (Fig. 1D–F), indicating that the prominent morphological abnormalities were mainly restricted to capitulum development rather than overall plant growth.

### Reiterative formation of secondary and tertiary floret-like organs in *marimo* capitula

To characterize the developmental features of the *marimo* phenotype, capitula were dissected and examined in detail under a stereomicroscope (Fig. 2). Secondary floret-like organs developed within primary floret-like organs, and tertiary floret-like organs were further formed within some of the secondary floret-like organs (Fig. 2A–D). These higher-order floret-like organs resembled the primary floret-like organs in morphology, and hierarchical and reiterative formation of floret-like organs occurred throughout the capitulum. Such secondary and tertiary floret-like organs were not observed in No. 251 or ‘Opal’.

**Figure 2.**
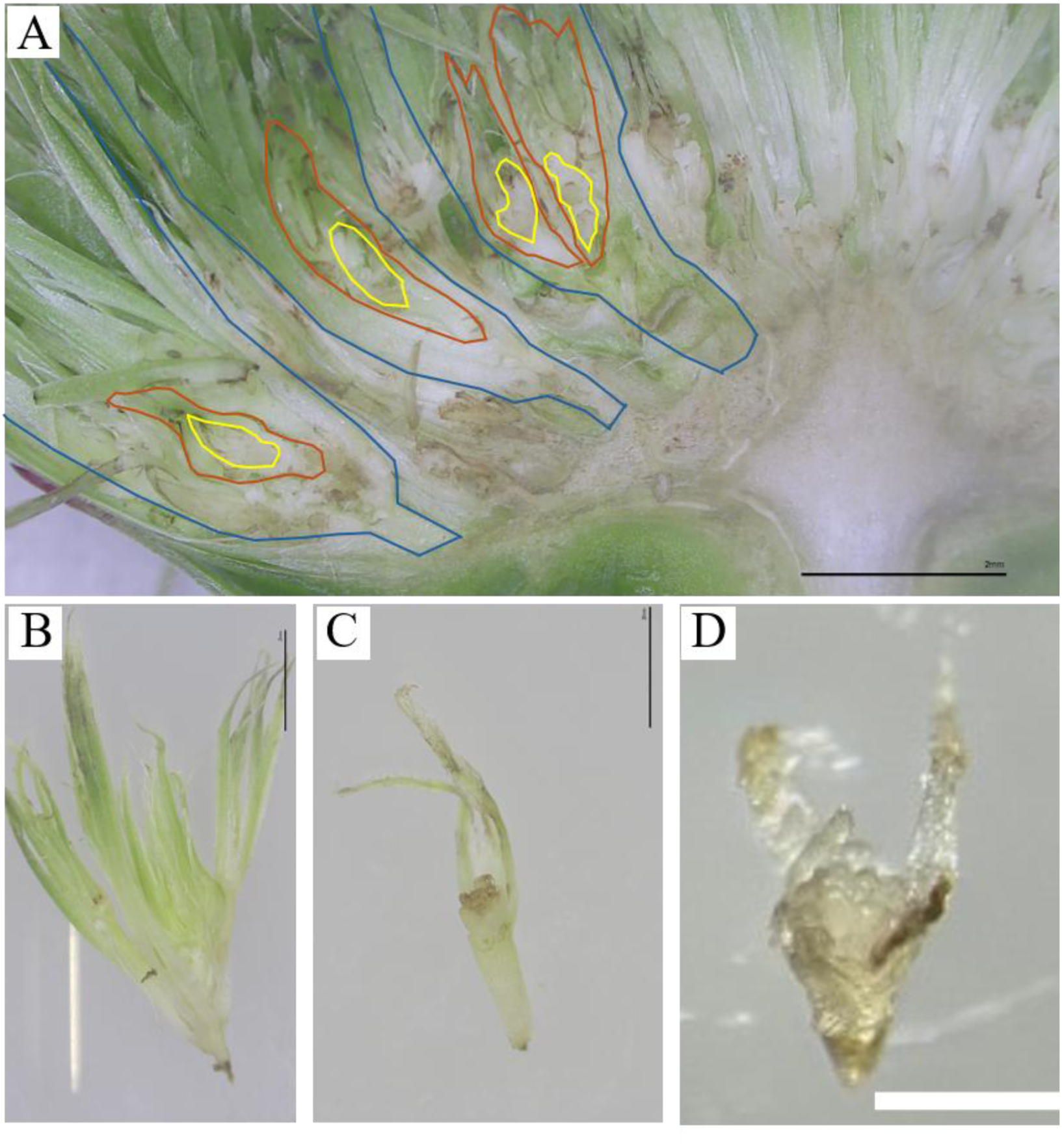
Reiterative formation of floret-like organs in the capitulum of the *marimo* mutant. (A) Longitudinal section of a capitulum of the *marimo* mutant. Primary, secondary, and tertiary floret-like organs are outlined in blue, orange, and yellow, respectively. (B) A primary floret-like organ isolated from the capitulum. (C) A secondary floret-like organ formed within a primary floret-like organ. (D) A tertiary floret-like organ formed within a secondary floret-like organ. Scale bars: A, 2 mm; B–D, 1 mm.

### Green organs of the *marimo* mutant exhibit epidermal features similar to those of involucral bracts

To characterize the green organs constituting the *marimo* capitulum, their epidermal structures were examined by scanning electron microscopy and compared with those of leaves, pappi, and involucral bracts from No. 251 and ‘Opal’ (Fig. 3). The epidermal cells of the elongated green organs in *marimo* were similar to those on the adaxial surfaces of involucral bracts in No. 251 and ‘Opal’, exhibiting elongated and curved shapes (Fig. 3A, B, H). By contrast, the pappi of No. 251 and ‘Opal’ consisted of numerous spine-like cells (Fig. 3D, G), whereas leaves displayed epidermal cell morphologies distinct from those of involucral bracts (Fig. 3C, F, I, J). Thus, the green organs constituting the *marimo* capitulum exhibited epidermal characteristics similar to those of wild-type involucral bracts.

**Figure 3.**
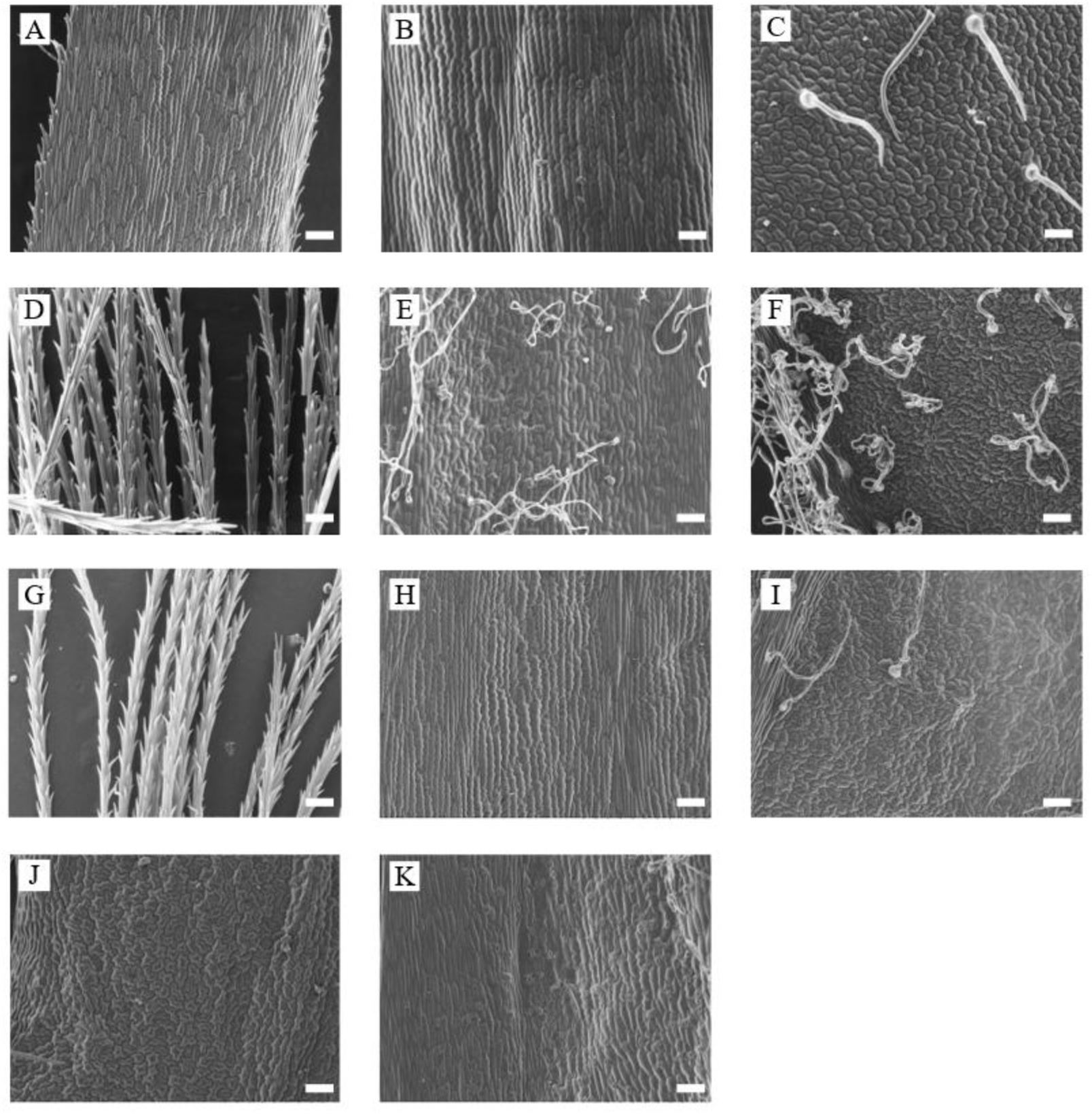
Scanning electron microscopy of green organs in the *marimo* mutant and involucral bracts, leaves, and pappi in wild-type gerbera. (A) A green organ constituting the capitulum of the *marimo* mutant. (B–F) Adaxial surface of an involucral bract (B), adaxial surface of a leaf (C), pappus (D), abaxial surface of an involucral bract (E), and abaxial surface of a leaf (F) in the wild-type cultivar ‘Opal’. (G–K) Pappus (G), adaxial surface of an involucral bract (H), abaxial surface of a leaf (I), adaxial surface of a leaf (J), and abaxial surface of an involucral bract (K) in the wild-type line No. 251. Scale bars: 100 µm.

### Extensive changes in gene expression occur in the *marimo* mutant

To investigate the molecular basis of the *marimo* phenotype, RNA-seq analysis was performed using developing capitula from the *marimo* mutant, No. 251, and ‘Opal’ (Fig. 4). Principal component analysis clearly separated the *marimo* samples from the No. 251 and ‘Opal’ samples, with the first principal component accounting for 81% of the total variance (Fig. 4A). Biological replicates of each genotype clustered closely together, indicating similar transcriptional profiles within each genotype. Differential expression analysis identified 1,981 significantly upregulated and 2,295 downregulated genes in *marimo* relative to ‘Opal’. In comparison with No. 251, 1,492 genes were upregulated and 2,296 were downregulated in *marimo*. Among these differentially expressed genes, 967 were commonly upregulated and 1,734 were commonly downregulated in both comparisons (Fig. 4B, C). These results demonstrated extensive transcriptional changes in developing *marimo* capitula relative to both wild types.

**Figure 4.**
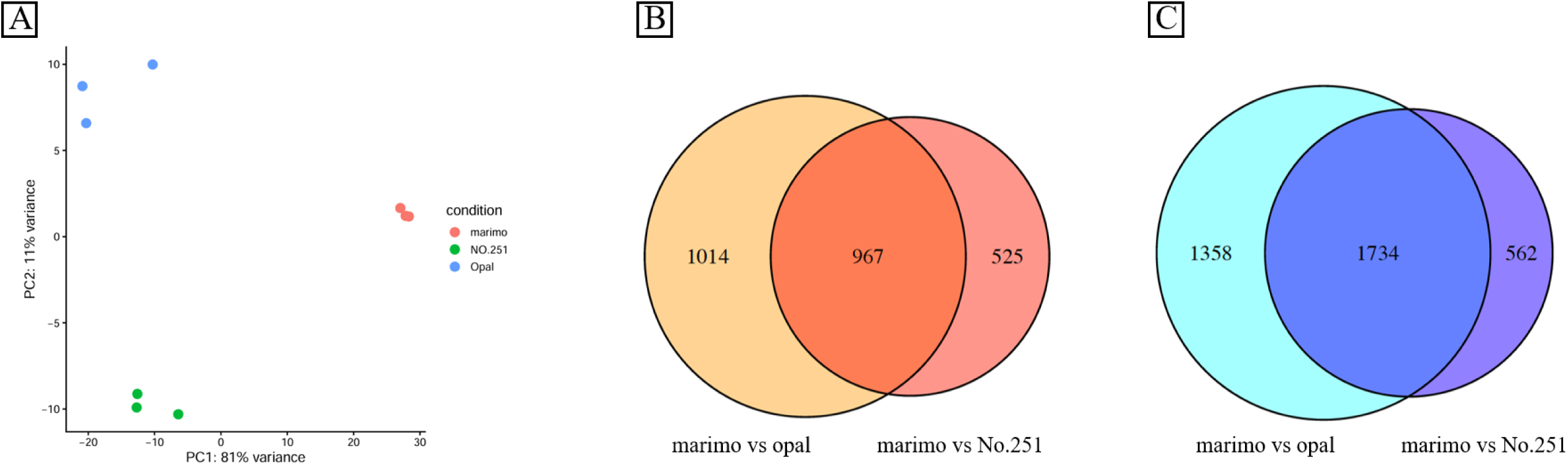
Transcriptome analysis of developing capitula from the *marimo* mutant and wild-type gerbera. (A) Principal component analysis of RNA-seq data obtained from developing capitula of the *marimo* mutant, the wild-type line No. 251, and the wild-type cultivar ‘Opal’. Principal components 1 and 2 account for 81% and 11% of the total variance, respectively. (B) Venn diagram showing the overlap of differentially expressed genes upregulated in the *marimo* mutant relative to ‘Opal’ and No. 251. (C) Venn diagram showing the overlap of differentially expressed genes downregulated in the *marimo* mutant relative to ‘Opal’ and No. 251.

### Protein–protein interaction network analysis identifies three major gene clusters associated with the marimo phenotype

To investigate functional relationships among genes associated with the marimo phenotype, a STRING-based protein–protein interaction network was constructed using Arabidopsis orthologs of the differentially expressed gerbera genes (Fig. 5). Markov clustering of the network identified three major clusters associated with tissue structure formation, reproductive organ development, and floral development (Fig. 5A–F).

**Figure 5.**
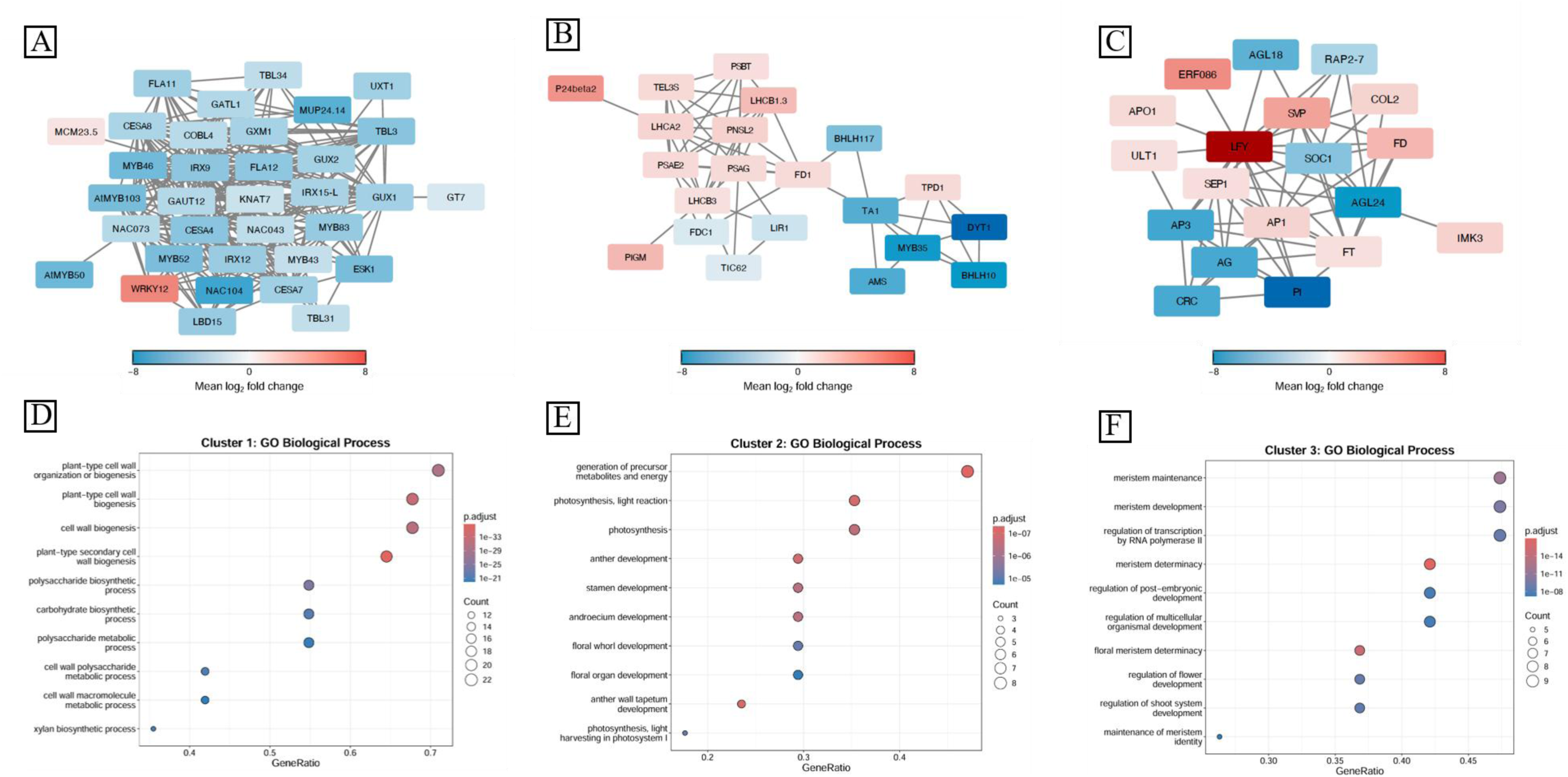
Protein–protein interaction network and Gene Ontology analyses of genes differentially expressed in the marimo mutant. (A–C) Protein–protein interaction networks constructed using Arabidopsis thaliana orthologs of genes differentially expressed in the marimo mutant and classified using the Markov Cluster Algorithm (MCL). Protein association information was obtained from the STRING database. Each node represents an Arabidopsis ortholog of a differentially expressed gerbera gene, and each edge represents a known or predicted protein–protein association retrieved from STRING. Node color indicates the mean log2 fold change across the comparisons of the marimo mutant with No. 251 and ‘Opal’. (A) MCL cluster 1. (B) MCL cluster 2. (C) MCL cluster 3. (D–F) Gene Ontology (GO) enrichment analyses of MCL cluster 1 (D), cluster 2 (E), and cluster 3 (F). The x-axis indicates the GeneRatio, and the y-axis indicates GO terms. Dot size represents the number of genes assigned to each GO term, and dot color indicates statistical significance based on the adjusted P value.

Cluster 1 contained genes associated with secondary cell wall formation and tissue structural development, including CESA4, CESA7, CESA8, MYB46, KNAT7, and members of the NAC transcription factor family (Fig. 5A, D).

Cluster 2 consisted of genes associated with stamen differentiation, anther development, and pollen formation, including AMS, DYT1, and MYB35 (Fig. 5B, E). Many of these genes were expressed at lower levels in *marimo* than in No. 251 and ‘Opal’.

Cluster 3 contained major regulators associated with floral organ identity and floral meristem development, including LFY, AP1, AG, AP3, the PISTILLATA/GLOBOSA-like B-class MADS-box gene GGLO1, and SEP1 (Fig. 5C). In this network, LFY was connected to a relatively large number of genes, whereas GGLO1 expression was markedly reduced in *marimo*.

Gene Ontology enrichment analysis revealed significant enrichment of biological processes associated with cell wall formation and organization in Cluster 1, stamen, anther, and male reproductive organ development in Cluster 2, and flower development, reproductive shoot system development, reproductive structure development, and meristem determinacy in Cluster 3 (Fig. 5D–F).

### Reduced GGLO1 expression is associated with the marimo phenotype

To validate the relationship between the reduced GGLO1 expression detected by RNA-seq and capitulum phenotype, GGLO1 expression was analyzed by RT-qPCR using a segregating population derived from self-pollination of No. 251 (Fig. S1). Before gene expression analysis, the progeny were classified as wild type or marimo type based on capitulum morphology. GGLO1 expression was consistently lower in the five marimo-type individuals than in the five wild-type individuals examined.

### Identification of a frameshift mutation in UFO

Because previous studies have shown that UFO plays an important role in gerbera capitulum development, the gene model corresponding to UFO was selected based on the annotation of the gerbera reference genome Ghyb_p1.1, and its coding sequence was analyzed. A single-nucleotide deletion affecting one of three consecutive cytosine residues was identified in the UFO coding region (ΔC) (Fig. 6A–C). The UFO coding sequence showed 99% nucleotide sequence identity across its full length with the previously reported GhUFO sequence (GenBank accession no. KU554695), confirming that the analyzed gene model corresponded to GhUFO. The UFO gene consisted of a single exon without introns.

**Figure 6.**
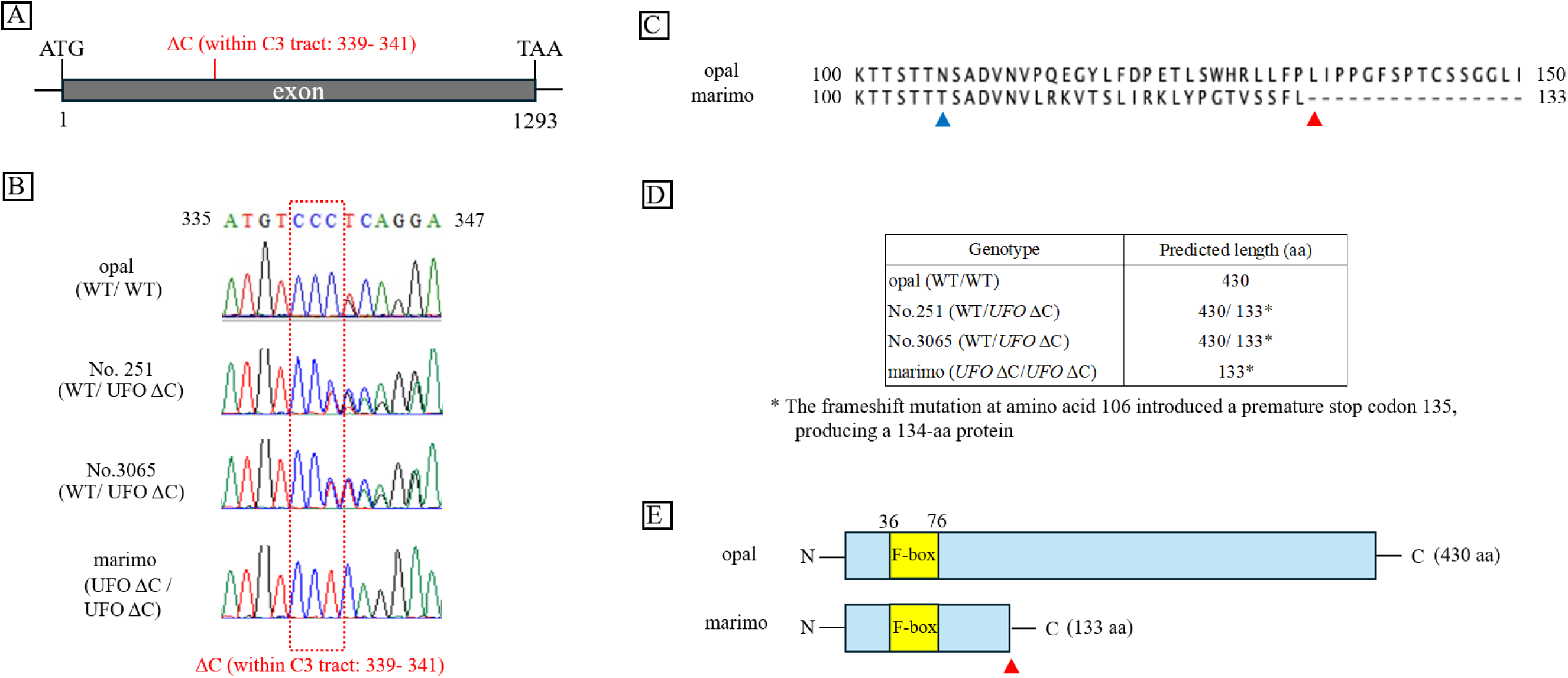
Identification of a frameshift mutation in the UNUSUAL FLORAL ORGANS gene of the *marimo* mutant. (A) Coding region of the UNUSUAL FLORAL ORGANS (*UFO*) gene model (Ghyb006_g00062.1) in the gerbera reference genome Ghyb_p1.1 and the position of the single-nucleotide deletion identified in the *marimo* mutant. The deletion is located within a stretch of three consecutive cytosine residues at positions 339–341 of the coding region. (B) Sequence chromatograms surrounding the mutation site in the wild-type cultivar ‘Opal’, the heterozygous lines No. 251 and No. 3065, and the *marimo* mutant. Red dashed boxes indicate the stretch of three consecutive cytosine residues and the position of the single-nucleotide deletion. (C) Alignment of the deduced amino acid sequences surrounding the mutation site in the wild type and the *marimo* mutant. The blue arrowhead indicates the start of the frameshift, and the red arrowhead indicates the premature termination site. (D) Predicted lengths of the UFO proteins encoded by each genotype. (E) Schematic representation of the UFO protein structures in the wild type and the *marimo* mutant. The yellow region indicates the F-box domain, and the red arrowhead indicates the termination site of the mutant protein.

The deletion was absent from the wild-type cultivar ‘Opal’ (WT/WT), whereas No. 251 and No. 3065 were heterozygous for the mutation (WT/UFOΔC), and the *marimo* mutant was homozygous for the mutant allele (UFOΔC/UFOΔC) (Fig. 6B, C). The presence of the mutant allele in the heterozygous state in both No. 251 and No. 3065 was consistent with the occurrence of *marimo*-type individuals among progeny from their reciprocal crosses.

The single-nucleotide deletion was predicted to alter the reading frame, substantially change the deduced amino acid sequence downstream of the mutation, and introduce a premature stop codon (Fig. 6D, E). Consequently, the mutant UFO protein was predicted to terminate after 134 amino acids and to be substantially shorter than the wild-type protein. Comparison of the deduced amino acid sequences confirmed divergence downstream of the mutation and premature termination of translation (Fig. S3). Although at least part of the N-terminal F-box domain was predicted to remain, most of the C-terminal region was lost. These results indicate that the single-nucleotide deletion in UFO is a strong candidate mutation associated with the *marimo* phenotype.

### The UFO mutation cosegregates with the *marimo* phenotype

To evaluate the genetic association between the UFO mutation and the *marimo* phenotype, segregation analysis was performed using 40 selfed progeny of No. 251 (Table 2). Genotyping identified nine wild-type homozygotes (WT/WT), 21 heterozygotes (WT/UFOΔC), and ten mutant homozygotes (UFOΔC/UFOΔC). This distribution did not differ significantly from the 1:2:1 ratio expected for segregation at a single locus (χ² = 0.150, P = 0.928). All ten mutant homozygotes exhibited the marimo phenotype, whereas all nine wild-type homozygotes and all 21 heterozygotes formed wild-type capitula. The resulting segregation of 30 wild-type and ten marimo-type individuals was fully consistent with the expected 3:1 ratio (χ² = 0.000, P = 1.000). No discordance between UFO genotype and capitulum phenotype was observed among the 40 individuals examined.

**Table 2.** Segregation of UFO genotypes and capitulum phenotypes among selfed progeny of No. 251 and No. 3065. Numbers of wild-type and *marimo*-type individuals according to UFO genotype among selfed progeny of No. 251 and No. 3065, both of which were heterozygous for the single-nucleotide deletion in *UFO*. Genotypes were classified as wild-type homozygous (WT/WT), heterozygous (WT/UFOΔC), or mutant homozygous (UFOΔC/UFOΔC).

| Genotype | No.251 S<sub>1</sub> progeny |  | No.3065 S<sub>1</sub> progeny |  |
| --- | --- | --- | --- | --- |
|  | WT | marimo | WT | marimo |
| WT/WT | 9 | 0 | 2 | 0 |
| WT/*UFO* ΔC | 21 | 0 | 4 | 0 |
| *UFO* ΔC/*UFO* ΔC | 0 | 10 | 0 | 2 |

Segregation analysis was also performed using eight selfed progeny of No. 3065. Two individuals were wild-type homozygotes, four were heterozygotes, and two were mutant homozygotes, exactly matching a 1:2:1 segregation ratio. Both mutant homozygotes exhibited the *marimo* phenotype, whereas all wild-type homozygotes and heterozygotes formed wild-type capitula. Together with the complete genotype–phenotype concordance observed among the 40 selfed progeny of No. 251, these results further support the association between homozygosity for the UFOΔC allele and the marimo phenotype.

## Discussion

### The *marimo* phenotype reflects defects in both floral organ identity and floral meristem determinacy

The *marimo* mutant identified in this study formed green, spherical, and compact capitula in which secondary and tertiary floret-like organs were reiteratively produced within primary floret-like organs (Figs. 1, 2). In addition, the epidermal structure of the green organs constituting the capitulum resembled that of wild-type involucral bracts (Fig. 3). These characteristics indicate that both floral organ identity and floral meristem determinacy are altered in the *marimo* mutant.

Altered function of floral organ identity genes can cause homeotic transformations in which one floral organ type is replaced by another (Bowman et al., 1991; Coen and Meyerowitz, 1991). In the *marimo* mutant, normal floral organs were replaced by green involucral bract-like organs, and new floret-like organs were formed within these structures. The *marimo* phenotype can therefore be regarded as a complex developmental abnormality involving both altered floral organ identity and reduced floral meristem determinacy.

A highly similar phenotype has been reported in transgenic gerbera plants in which GhUFO expression was suppressed by RNA interference. In these plants, normal floral organs were replaced by green involucral bract-like organs, and secondary and tertiary floret-like organs were reiteratively formed (Zhao et al., 2016). By analyzing a naturally occurring frameshift mutant of UFO and its segregating progeny, the present study provides new genetic evidence that altered UFO function affects both floral organ identity and floral meristem determinacy.

### The UFO frameshift mutation is a strong candidate underlying the *marimo* phenotype

A single-nucleotide deletion causing a frameshift was identified in the coding region of UFO (Fig. 6). This mutation was predicted to alter the deduced amino acid sequence downstream of the deletion and introduce a premature stop codon. Consequently, the mutant protein was predicted to terminate after 134 amino acids, resulting in the loss of most of the C-terminal region of the wild-type UFO protein (Fig. 6; Fig. S3). This extensive change in the predicted protein structure suggests that the function of the mutant UFO protein is likely to be severely impaired.

In selfed progeny of No. 251 and No. 3065, only UFOΔC/UFOΔC individuals exhibited the *marimo* phenotype, whereas WT/WT and WT/UFOΔC individuals showed the wild-type phenotype. UFO genotype and capitulum phenotype were fully concordant among all individuals examined (Table 2). The UFOΔC allele was present in the heterozygous state in both No. 251 and No. 3065, which produced *marimo*-type progeny, and in the homozygous state in the original *marimo* mutant. In reciprocal crosses between No. 251 and No. 3065, *marimo*-type and wild-type individuals segregated at an approximate ratio of 1:3 in both crossing directions. These findings are consistent with a genetic model in which both lines carry the same recessive mutant allele in the heterozygous state and the *marimo* phenotype appears when the UFOΔC allele becomes homozygous in the progeny. The occurrence of the phenotype in both reciprocal crosses also supports inheritance through a recessive mutation in the nuclear genome.

Taken together with the close morphological similarity between the *marimo* mutant and GhUFO RNAi plants (Zhao et al., 2016), these genetic results indicate that the UFO frameshift mutation is the strongest candidate variant underlying the *marimo* phenotype. Complementation with a wild-type UFO allele and recreation of the same mutation by genome editing will further establish the causal relationship between the mutation and the phenotype.

### Changes in the expression of floral development-related genes associated with the UFO mutation

RNA-seq analysis revealed extensive differences in gene expression between the *marimo* mutant and the wild types (Fig. 4). In the principal component analysis, *marimo* samples were clearly separated from No. 251 and ‘Opal’, and the first principal component accounted for 81% of the total variance. These results indicate that developing *marimo* capitula have a markedly different transcriptional state from those of the wild types.

STRING-based protein–protein interaction network analysis of Arabidopsis orthologs identified three major clusters associated with tissue structure formation, reproductive organ differentiation, and floral organ development (Fig. 5). Cluster 3 contained major regulators of floral organ identity and floral meristem development, including LFY, AP1, AG, AP3, GGLO1, and SEP1.

RNA-seq analysis showed increased LFY expression and markedly reduced expression of GGLO1, a PISTILLATA/GLOBOSA-like B-class MADS-box gene, in the marimo mutant. RT-qPCR analysis of selfed progeny of No. 251 further confirmed that GGLO1 expression was consistently lower in marimo-type individuals than in wild-type individuals. In *Arabidopsis*, UFO acts together with LFY to activate the B-class floral organ identity genes AP3 and PI (Lee et al., 1997; Zhao et al., 2001; Chae et al., 2008). UFO has also been shown to form a complex with LFY and DNA and to alter the target sequence recognition of LFY (Rieu et al., 2023). These findings suggest that the reduced GGLO1 expression in *marimo* is associated not with insufficient LFY expression, but with altered LFY–UFO-dependent transcriptional regulation caused by impaired UFO function.

The increased LFY expression in *marimo* may reflect an increased proportion of LFY-expressing meristematic tissues resulting from the reiterative formation of secondary and tertiary floret-like organs. In gerbera, GhLFY is expressed in young capitulum meristems and developing floret primordia and contributes to capitulum determinacy and floret formation (Zhao et al., 2016). Increased LFY expression may also represent a feedback response to altered progression of the floral developmental program.

The RNA-seq results showed increased LFY expression in the marimo mutant despite its severe floral developmental abnormalities, supporting the importance of functional cooperation between LFY and UFO in gerbera capitulum development. Analysis of LFY- and UFO-binding regions and transcriptional assays using the GGLO1 promoter will help determine whether GGLO1 is a direct target of the LFY–UFO regulatory module.

### Reduced expression of GGLO1 and other floral development genes is associated with the formation of involucral bract-like organs

In the *marimo* mutant, normal floral organs were replaced by elongated green organs whose epidermal structure resembled that of wild-type involucral bracts (Fig. 3). This finding suggests that floral organ identity was not properly established and that the resulting organs acquired involucral bract-like or vegetative characteristics.

Transcriptome analysis revealed reduced expression of AMS and DYT1, which are involved in tapetum, anther, and pollen development, as well as reduced expression of MYB35 and the B-class floral organ identity gene GGLO1 (Sorensen et al., 2003; Zhang et al., 2006). GGLO1 is mainly expressed in petals and stamens, and its suppression causes homeotic transformation of these organs, indicating that it plays an important role in establishing petal and stamen identity in gerbera (Broholm et al., 2010).

Reduced expression of GGLO1 and other genes involved in floral and reproductive organ development may therefore have contributed to defective petal and stamen formation and the conversion of normal floral organs into green involucral bract-like organs in the *marimo* mutant. This interpretation is consistent with the conversion of normal floral organs into involucral bract-like organs in GhUFO RNAi plants (Zhao et al., 2016). By contrast, the reiterative formation of secondary and tertiary floret-like organs is more likely associated with altered floral meristem identity and determinacy resulting from impaired UFO function.

### Morphological variation among *marimo*-type progeny

Morphological variation in involucral bract-like organs was observed among marimo-type progeny. Representative variation among selfed progeny of No. 251 is shown in Fig. 7. In some individuals, these organs were markedly elongated, whereas in others they were extremely short, giving the capitulum surface a bristle-like appearance. In addition to individuals producing green organs, some individuals formed organs with reddish pigmentation.

**Figure 7.**
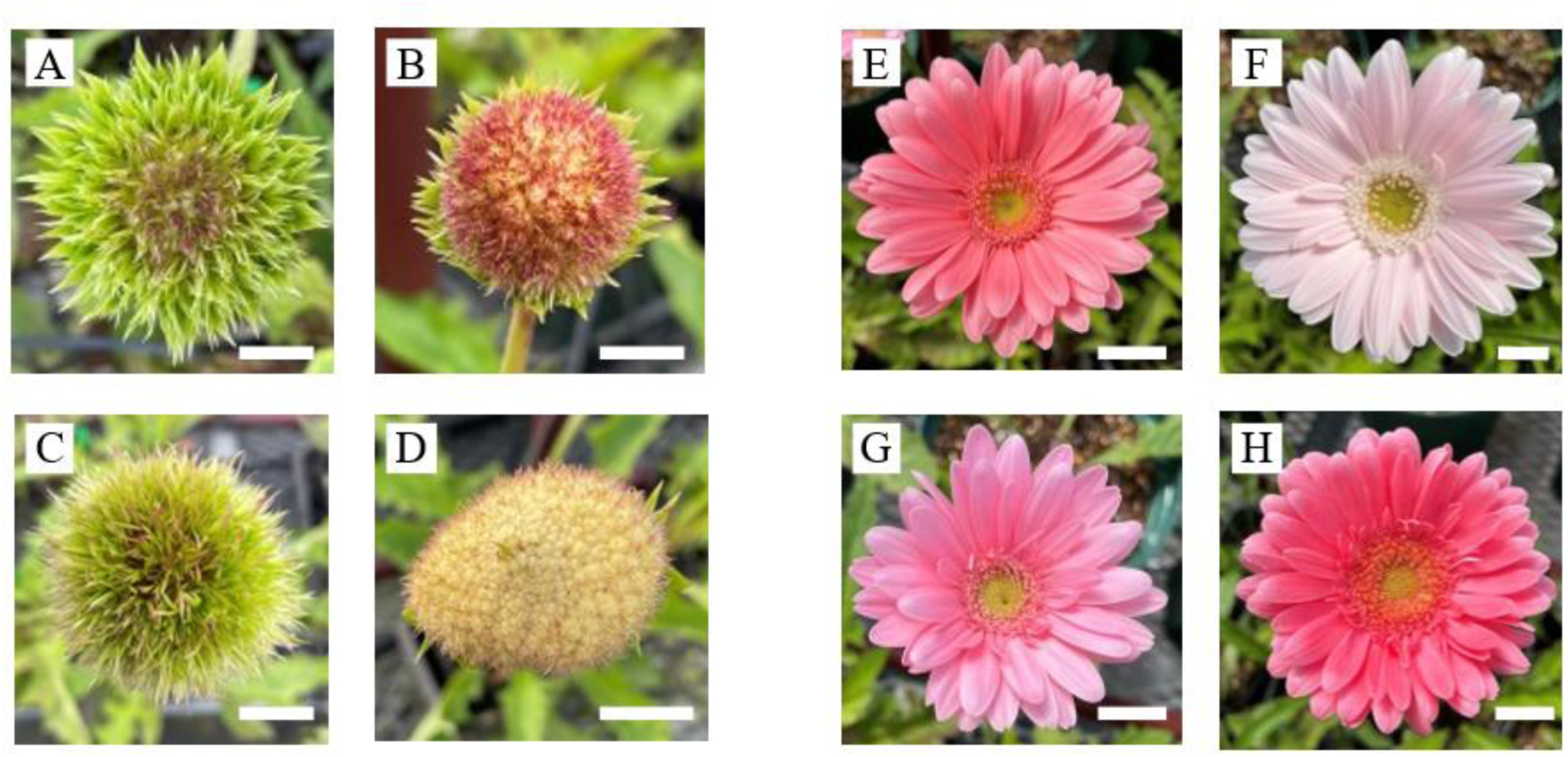
Representative *marimo*-type and wild-type capitula among selfed progeny of No. 251. Representative capitula among selfed progeny of No. 251. (A–D) *Marimo*-type capitula. (E–H) Wild-type capitula. Scale bars: 1 cm.

All of these individuals shared the fundamental features of the *marimo* phenotype, namely defective formation of normal floral organs and reiterative production of floret-like organs. These observations suggest that homozygosity for the UFOΔC allele alters floral organ identity and floral meristem determinacy, thereby specifying the basic architecture of the *marimo*-type capitulum. By contrast, the length, degree of development, and coloration of the involucral bract-like organs may be influenced by additional genetic factors and differences in genetic background.

Quantification of involucral bract-like organ length, number, pigmentation, and overall capitulum morphology, followed by genetic analysis using segregating populations, should facilitate the identification of modifier genes controlling these traits. In particular, comparison of individuals that are homozygous for UFOΔC but differ in organ length or coloration may enable efficient identification of genetic factors regulating organ morphology within a shared developmental background.

In the present study, these organs were classified as involucral bract-like based on the similarity of their epidermal structure to that of normal involucral bracts. Histological observations at early developmental stages and expression analysis of genes associated with involucral bract development will help clarify the developmental relationship between these organs and normal involucral bracts. The *marimo* mutant and its segregating populations are therefore expected to provide useful genetic materials for investigating the regulation of involucral bract-like organ length, morphology, and coloration in Asteraceae.

### Changes in the expression of cell wall-related genes reflect altered tissue composition of the capitulum

Network analysis identified a group of genes associated with secondary cell wall formation and cell wall organization, including CESA4, CESA7, CESA8, MYB46, KNAT7, and several NAC transcription factors (Fig. 5). In *Arabidopsis*, CESA4, CESA7, and CESA8 are major components of the cellulose synthase complex involved in secondary cell wall formation, whereas MYB46, KNAT7, and NAC transcription factors participate in the transcriptional regulation of secondary cell wall-related genes (Zhong et al., 2008; Kim et al., 2013).

Because whole developing capitula were used for RNA-seq analysis, the observed expression levels likely reflected not only changes in gene expression within individual tissues but also differences in the types and relative proportions of tissues constituting the capitulum. In the *marimo* mutant, petal and reproductive organ development was strongly suppressed, and most of the capitulum consisted of green involucral bract-like organs. Therefore, the altered expression of cell wall-related genes likely reflects changes in the overall tissue composition of the capitulum associated with the replacement of floral tissues, including petals and reproductive organs, by involucral bract-like tissues, in addition to possible changes in cell wall formation within individual organs.

This interpretation is consistent with the morphological observation that the epidermal structure of the green organs in *marimo* resembled that of wild-type involucral bracts. Separate collection and comparison of wild-type petals, wild-type involucral bracts, and *marimo* involucral bract-like organs for gene expression and cell wall composition analyses will help determine whether the observed changes in cell wall-related gene expression primarily result from altered tissue composition or reflect specific cell wall characteristics of the involucral bract-like organs.

### Developmental model and breeding potential of the *marimo* phenotype

Based on the present results, LFY and UFO are likely to function cooperatively in wild-type gerbera to regulate the expression of floral organ development-related genes, including GGLO1, and to support normal floral meristem development. In the *marimo* mutant, the UFO frameshift mutation may reduce the activity of the LFY–UFO regulatory module, thereby altering both the transcriptional regulation of floral organ development-related genes and floral meristem determinacy (Fig. 8). This model is consistent with findings in *Arabidopsis* showing that LFY and UFO cooperatively regulate B-class gene transcription (Lee et al., 1997; Zhao et al., 2001; Chae et al., 2008; Rieu et al., 2023), as well as with the phenotype reported in GhUFO RNAi gerbera plants (Zhao et al., 2016).

**Figure 8.**
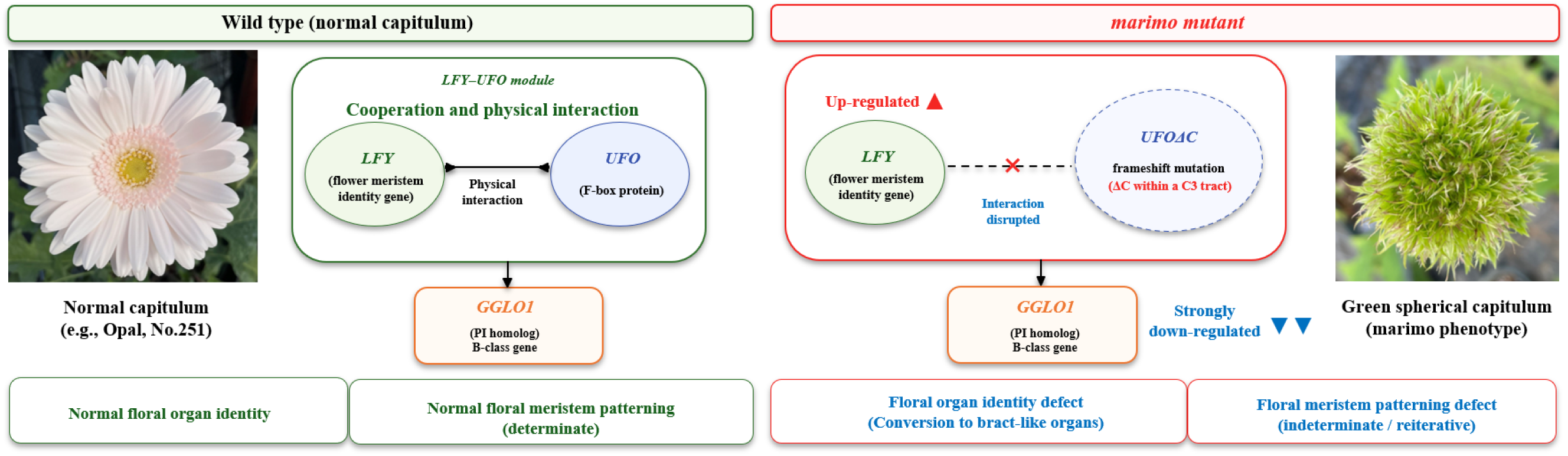
Proposed developmental model linking the UFO frameshift mutation to the marimo phenotype. Schematic model of the LFY–UFO regulatory module, expression of floral organ development-related genes including GGLO1, floral organ identity, and floral meristem determinacy in wild-type gerbera and the *marimo* mutant. The left panel shows normal capitulum development in the wild type, whereas the right panel shows the formation of a green, spherical capitulum in the *marimo* mutant carrying the frameshift mutation in *UFO*. Solid arrows indicate known or inferred regulatory relationships, whereas dashed arrows indicate regulatory relationships that have not been directly demonstrated in gerbera. The cross indicates the predicted reduction in LFY–UFO regulatory activity associated with the truncated UFO protein.

Although RNA-seq analysis showed increased LFY expression in the marimo mutant, GGLO1 expression was markedly reduced. Altered LFY–UFO-dependent transcriptional regulation resulting from impaired UFO function may have changed the expression of GGLO1 and other floral organ development-related genes, thereby preventing the proper establishment of floral organ identity. By contrast, the reiterative production of secondary and tertiary floret-like organs likely resulted from altered floral meristem identity and determinacy caused by reduced UFO function.

The UFOΔC allele appears to specify the basic architecture of the *marimo*-type capitulum, whereas the length and coloration of the involucral bract-like organs may be influenced by other genetic factors. The *marimo* mutant and its segregating populations are therefore expected to serve as valuable genetic materials for investigating not only the regulation of floral organ development and floral meristem determinacy by the LFY–UFO module, but also the molecular mechanisms controlling the morphology and coloration of involucral bract-like organs.

The spherical and compact capitulum of the *marimo* mutant also represents a distinctive ornamental trait that differs clearly from those of conventional gerbera cultivars. DNA markers based on the UFOΔC allele could enable the selection of phenotypically normal heterozygous individuals at the seedling stage. Furthermore, combining the UFOΔC allele with diverse genetic backgrounds may facilitate the development of a range of *marimo*-type cultivars differing in the length and coloration of their involucral bract-like organs.

Future complementation and genome-editing experiments will allow the causal relationship between the UFO mutation and the *marimo* phenotype to be tested directly. Analysis of GGLO1 transcriptional regulation by the LFY–UFO complex will further clarify the molecular basis of the phenotype. In addition, identifying modifier genes controlling the length, degree of development, and coloration of involucral bract-like organs using *marimo* segregating populations will advance our understanding of the genetic basis underlying complex capitulum development and ornamental trait diversity in Asteraceae.

## Conclusions

In this study, we identified a novel gerbera mutant, *marimo*, characterized by green, spherical, and compact capitula, and investigated its morphological, transcriptomic, and genetic features. In the *marimo* mutant, green involucral bract-like organs developed at positions normally occupied by floral organs, and secondary and tertiary floret-like organs were reiteratively formed within primary floret-like organs. These characteristics suggest that the *marimo* phenotype involves both altered floral organ identity and reduced floral meristem determinacy.

RNA-seq analysis revealed extensive changes in the expression of genes associated with floral organ development, reproductive organ differentiation, and tissue composition. In addition, a single-nucleotide deletion causing a frameshift was identified in the coding region of UFO. This mutation was present in the heterozygous state in No. 251 and No. 3065 and in the homozygous state in the *marimo* mutant. In selfed progeny of both lines, UFO genotype was fully concordant with capitulum phenotype, whereas reciprocal crosses between No. 251 and No. 3065 produced *marimo*-type and wild-type progeny at an approximate ratio of 1:3.

Taken together, these results identify the naturally occurring frameshift mutation in UFO as the strongest candidate variant associated with the marimo phenotype. Reduced activity of the LFY–UFO regulatory module associated with this mutation may have affected both the expression of floral organ development-related genes and floral meristem determinacy, thereby giving rise to the distinctive capitulum architecture of *marimo*.

Morphological variation in the length and coloration of involucral bract-like organs was also observed among *marimo*-type progeny, suggesting that genetic factors other than UFOΔC contribute to these traits. The *marimo* mutant and its segregating populations therefore provide useful genetic resources for investigating floral organ development, floral meristem determinacy, and involucral bract-like organ morphogenesis in Asteraceae. They may also serve as valuable breeding materials for introducing novel ornamental traits into gerbera.

## Availability of data and materials

The RNA-seq data generated in this study have been deposited in the DDBJ Sequence Read Archive under BioProject accession number PRJDB35793. The BioSample accession numbers for the marimo mutant and No. 251 samples are SAMD01948570–SAMD01948575. The corresponding DRA Run accession numbers will be added upon completion of the DRA submission.

---

**Figure S1.**
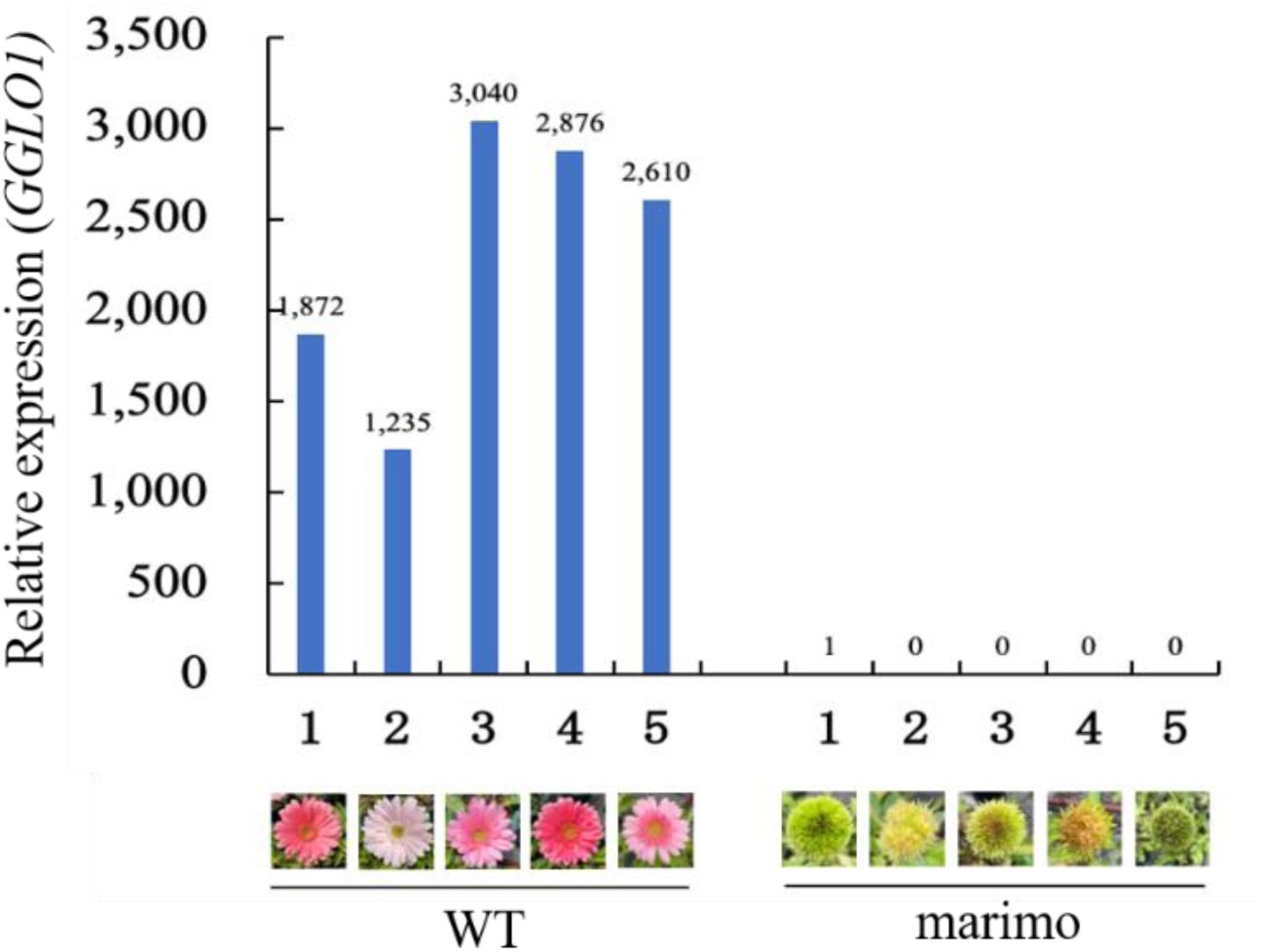
Relative expression levels of GGLO1 among selfed progeny of No. 251. Relative expression levels of GGLO1 in five selfed progeny of No. 251 that formed wild-type capitula and five that formed *marimo*-type capitula. The capitulum of each individual analyzed is shown below the corresponding bar. Expression levels were normalized using 2PS as the reference gene.

**Figure S2.**
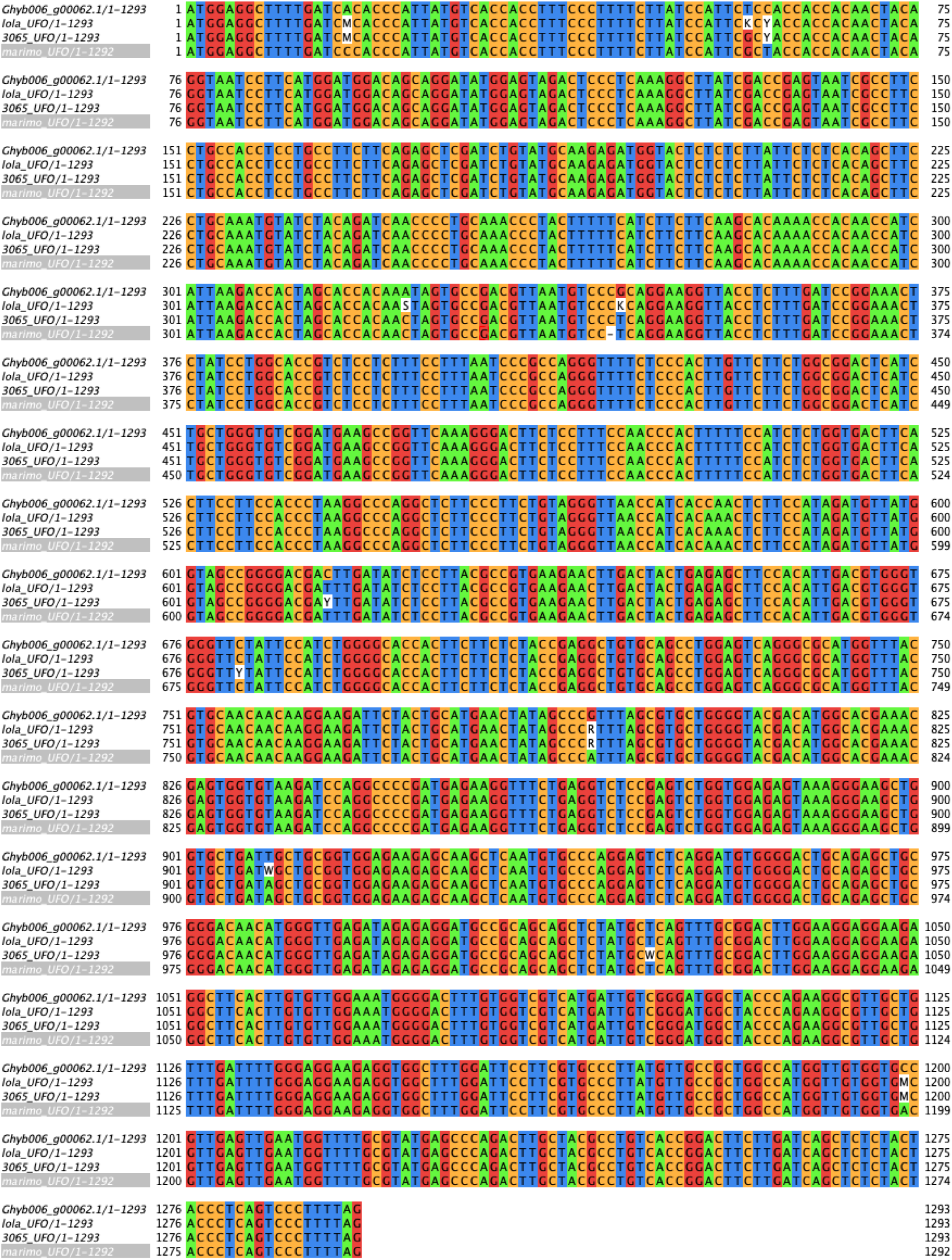
Nucleotide sequence alignment of the UFO coding region in wild-type and UFO mutant lines. Alignment of the coding-region nucleotide sequences of the *UFO* gene model in the gerbera reference genome Ghyb_p1.1 and those obtained from the wild-type cultivar ‘Opal’, No. 251, No. 3065, and the *marimo* mutant. Nucleotide positions are shown at the right end of each row.

**Figure S3.**
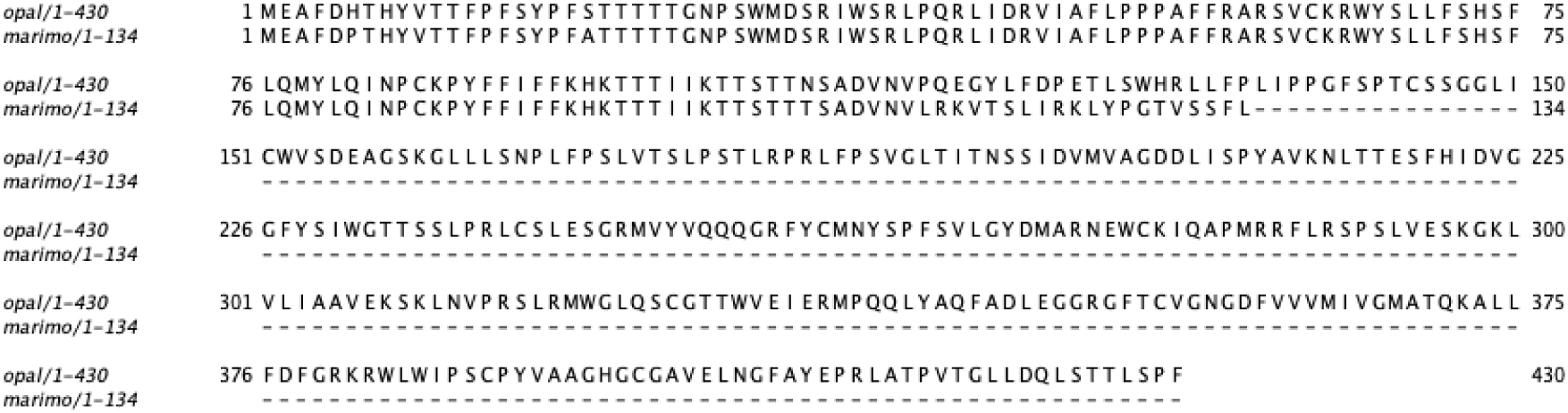
Alignment of the deduced UFO amino acid sequences in the wild type and the *marimo* mutant. Alignment of the deduced UFO amino acid sequences from the wild-type cultivar ‘Opal’ and the *marimo* mutant. Amino acid positions are shown at the right end of each row. Hyphens indicate regions in which no corresponding amino acid sequence is present.

## References

Ai Y, Zhang C, Sun Y, Wang W, He Y, Bao M. Characterization and functional analysis of five MADS-box B class genes related to floral organ identification in Tagetes erecta. PLoS One. 2017;12:e0169777. 10.1371/journal.pone.0169777

Andrews S. FastQC: a quality control tool for high throughput sequence data. Cambridge: Babraham Bioinformatics, Babraham Institute; 2010. Available from: https://www.bioinformatics.babraham.ac.uk/projects/fastqc/

Aoyagi YB, Shimada R, Hirakawa H, Toyoda A, Toh H, Isobe S, et al. Chromosome-level genome assembly of the Gerbera (Gerbera hybrida) using HiFi long-read and Hi-C technologies. DNA Res. 2026;33:dsag002. 10.1093/dnares/dsag002

Bolger AM, Lohse M, Usadel B. Trimmomatic: a flexible trimmer for Illumina sequence data. Bioinformatics. 2014;30:2114–2120. 10.1093/bioinformatics/btu170

Bowman JL, Smyth DR, Meyerowitz EM. Genetic interactions among floral homeotic genes of Arabidopsis. Development. 1991;112:1–20. 10.1242/dev.112.1.1

Broholm SK, Pöllänen E, Ruokolainen S, Tähtiharju S, Kotilainen M, Albert VA, et al. Functional characterization of B class MADS-box transcription factors in Gerbera hybrida. J Exp Bot. 2010;61:75–85. 10.1093/jxb/erp279

Broholm SK, Teeri TH, Elomaa P. Molecular control of inflorescence development in Asteraceae. Adv Bot Res. 2014;72:297–333. 10.1016/B978-0-12-417162-6.00010-9

Buchfink B, Reuter K, Drost HG. Sensitive protein alignments at tree-of-life scale using DIAMOND. Nat Methods. 2021;18:366–368. 10.1038/s41592-021-01101-x

Chae E, Tan QKG, Hill TA, Irish VF. An Arabidopsis F-box protein acts as a transcriptional co-factor to regulate floral development. Development. 2008;135:1235–1245. 10.1242/dev.015842

Coen ES, Meyerowitz EM. The war of the whorls: genetic interactions controlling flower development. Nature. 1991;353:31–37. 10.1038/353031a0

Ditta G, Pinyopich A, Robles P, Pelaz S, Yanofsky MF. The SEP4 gene of Arabidopsis thaliana functions in floral organ and meristem identity. Curr Biol. 2004;14:1935–1940. 10.1016/j.cub.2004.10.028

Dong ZC, Zhao Z, Liu CW, Luo JH, Yang J, Huang WH, et al. Floral patterning in Lotus japonicus. Plant Physiol. 2005;137:1272–1282. 10.1104/pp.104.054288

Elomaa P, Zhao Y, Zhang T. Flower heads in Asteraceae—recruitment of conserved developmental regulators to control the flower-like inflorescence architecture. Hortic Res. 2018;5:36. 10.1038/s41438-018-0056-8

Ferguson AC, Pearce S, Band LR, Rodriguez-Enriquez J, Hodgman TC, Murchie EH, et al. BHLH10, BHLH89 and BHLH91 are redundant regulators of tapetum development and reproductive fate in Arabidopsis. Plant Physiol. 2017;175:1162–1177. 10.1104/pp.17.00514

Fu Y, Esselink GD, Visser RGF, van Tuyl JM, Arens P. Transcriptome analysis of Gerbera hybrida including in silico confirmation of defense genes found. Front Plant Sci. 2016;7:247. 10.3389/fpls.2016.00247

Funk VA, Susanna A, Stuessy TF, Bayer RJ, editors. Systematics, evolution, and biogeography of Compositae. Vienna: International Association for Plant Taxonomy; 2009. DOI not assigned.

Gurung V, Muñoz-Gómez S, Jones DS. Putting heads together: developmental genetics of the Asteraceae capitulum. Curr Opin Plant Biol. 2024;79:102589. 10.1016/j.pbi.2024.102589

Hepworth SR, Klenz JE, Haughn GW. UFO in the Arabidopsis inflorescence apex is required for floral-meristem identity and bract suppression. Planta. 2006;223:769–778. 10.1007/s00425-005-0138-3

Kim D, Paggi JM, Park C, Bennett C, Salzberg SL. Graph-based genome alignment and genotyping with HISAT2 and HISAT-genotype. Nat Biotechnol. 2019;37:907–915. 10.1038/s41587-019-0201-4

Kim M, Cui ML, Cubas P, Gillies A, Lee K, Chapman MA, et al. Regulatory genes control a key morphological and ecological trait transferred between species. Science. 2008;322:1116– 1119. 10.1126/science.1164371

Kim WC, Ko JH, Kim JY, Kim J, Bae HJ, Han KH. MYB46 directly regulates the gene expression of secondary wall-associated cellulose synthases in Arabidopsis. Plant J. 2013;73:26–36. 10.1111/j.1365-313X.2012.05124.x

Kloos WE, George L, Suttors S. Inheritance of flower doubleness and laciniation in spider gerberas. J Am Soc Hortic Sci. 2004;129:802–810. 10.21273/JASHS.129.6.0802

Lee I, Wolfe DS, Nilsson O, Weigel D. A LEAFY co-regulator encoded by UNUSUAL FLORAL ORGANS. Curr Biol. 1997;7:95–104. 10.1016/S0960-9822(06)00053-4

Levin JZ, Meyerowitz EM. UFO: an Arabidopsis gene involved in both floral meristem and floral organ development. Plant Cell. 1995;7:529–548. 10.1105/tpc.7.5.529

Liao Y, Smyth GK, Shi W. featureCounts: an efficient general-purpose program for assigning sequence reads to genomic features. Bioinformatics. 2014;30:923–930. 10.1093/bioinformatics/btt656

Love MI, Huber W, Anders S. Moderated estimation of fold change and dispersion for RNA-seq data with DESeq2. Genome Biol. 2014;15:550. 10.1186/s13059-014-0550-8

Naing AH, Lee JH, Park KI, Kim CK. Transcriptional control of anthocyanin biosynthesis genes and transcription factors associated with flower coloration patterns in *Gerbera hybrida*. 3 Biotech. 2018;8:65. 10.1007/s13205-018-1099-0

Nakano M, Hirakawa H, Fukai E, Toyoda A, Kajitani R, Minakuchi Y, et al. A chromosome-level genome sequence of Chrysanthemum seticuspe, a model species for hexaploid cultivated chrysanthemum. Commun Biol. 2021;4:1167. 10.1038/s42003-021-02704-y

Pelaz S, Ditta GS, Baumann E, Wisman E, Yanofsky MF. B and C floral organ identity functions require SEPALLATA MADS-box genes. Nature. 2000;405:200–203. 10.1038/35012103

R Core Team. R: a language and environment for statistical computing. Vienna: R Foundation for Statistical Computing; 2025. Available from: https://www.R-project.org/

Ratcliffe OJ, Bradley DJ, Coen ES. Separation of shoot and floral identity in Arabidopsis. Development. 1999;126:1109–1120. 10.1242/dev.126.6.1109

Rieu P, Turchi L, Thévenon E, Zarkadas E, Nanao M, Chahtane H, et al. The F-box protein UFO controls flower development by redirecting the master transcription factor LEAFY to new cis-elements. Nat Plants. 2023;9:315–329. 10.1038/s41477-023-01336-y

Samach A, Klenz JE, Kohalmi SE, Risseeuw E, Haughn GW, Crosby WL. The UNUSUAL FLORAL ORGANS gene of Arabidopsis thaliana is an F-box protein. Plant J. 1999;20:433–445. 10.1046/j.1365-313X.1999.00617.x

Shannon P, Markiel A, Ozier O, Baliga NS, Wang JT, Ramage D, et al. Cytoscape: a software environment for integrated models of biomolecular interaction networks. Genome Res. 2003;13:2498–2504. 10.1101/gr.1239303

Sorensen AM, Kröber S, Unte US, Huijser P, Dekker K, Saedler H. The ABORTED MICROSPORES (AMS) gene encodes a MYC class transcription factor. Plant J. 2003;33:413–423. 10.1046/j.1365-313X.2003.01644.x

Szklarczyk D, Kirsch R, Koutrouli M, Nastou K, Mehryary F, Hachilif R, et al. The STRING database in 2025: protein networks with directionality of regulation. Nucleic Acids Res. 2025;53:D730–D737. 10.1093/nar/gkae1113

Uimari A, Kotilainen M, Elomaa P, Yu D, Albert VA, Teeri TH. Integration of reproductive meristem fates by a SEPALLATA-like MADS-box gene. Proc Natl Acad Sci U S A. 2004;101:15817–15822. 10.1073/pnas.0405310101

Utriainen M, Morris JH. clusterMaker2: a major update to clusterMaker, a multi-algorithm clustering app for Cytoscape. BMC Bioinformatics. 2023;24:134. 10.1186/s12859-023-05225-z

Weigel D, Alvarez J, Smyth DR, Yanofsky MF, Meyerowitz EM. LEAFY controls floral meristem identity in Arabidopsis. Cell. 1992;69:843–859. 10.1016/0092-8674(92)90295-N

Weigel D, Meyerowitz EM. The ABCs of floral homeotic genes. Cell. 1994;78:203–209. 10.1016/0092-8674(94)90291-7

Weigel D, Nilsson O. A developmental switch sufficient for flower initiation in diverse plants. Nature. 1995;377:495–500. 10.1038/377495a0

Wickham H. ggplot2: elegant graphics for data analysis. New York: Springer-Verlag; 2016. 10.1007/978-3-319-24277-4

Wilkinson MD, Haughn GW. UNUSUAL FLORAL ORGANS controls meristem identity and organ primordia fate in Arabidopsis. Plant Cell. 1995;7:1485–1499. 10.1105/tpc.7.9.1485

Wu T, Hu E, Xu S, Chen M, Guo P, Dai Z, et al. clusterProfiler 4.0: a universal enrichment tool for interpreting omics data. Innovation (Camb). 2021;2:100141. 10.1016/j.xinn.2021.100141

Xia Y, Shi M, Chen W, Hu X, Zhang L, Jiang Y, et al. Expression pattern and functional characterization of PISTILLATA ortholog in Eriobotrya japonica. Front Plant Sci. 2020;10:1685. 10.3389/fpls.2019.01685

Yu D, Kotilainen M, Pöllänen E, Mehto M, Elomaa P, Helariutta Y, et al. Organ identity genes and modified patterns of flower development in Gerbera hybrida. Plant J. 1999;17:51–62. 10.1046/j.1365-313X.1999.00351.x

Zhang T, Zhao Y, Juntheikki I, Mouhu K, Broholm SK, Rijpkema AS, et al. Dissecting functions of SEPALLATA-like MADS-box genes in patterning of the pseudanthial inflorescence of gerbera. New Phytol. 2017;216:939–954. 10.1111/nph.14707

Zhang T, Cieslak M, Owens A, Wang F, Broholm SK, Teeri TH, et al. Phyllotactic patterning of gerbera flower heads. Proc Natl Acad Sci U S A. 2021;118:e2016304118. 10.1073/pnas.2016304118

Zhang T, Elomaa P. Development and evolution of the Asteraceae capitulum. New Phytol. 2024;242:33–48. 10.1111/nph.19554

Zhang W, Sun Y, Timofejeva L, Chen C, Grossniklaus U, Ma H. Regulation of Arabidopsis tapetum development and function by DYSFUNCTIONAL TAPETUM1 (DYT1) encoding a putative bHLH transcription factor. Development. 2006;133:3085–3095. 10.1242/dev.02463

Zhao D, Yu Q, Chen M, Ma H. The ASK1 gene regulates B function gene expression in cooperation with UFO and LEAFY in Arabidopsis. Development. 2001;128:2735–2746. 10.1242/dev.128.14.2735

Zhao Y, Zhang T, Broholm SK, Tähtiharju S, Mouhu K, Albert VA, et al. Evolutionary co-option of floral meristem identity genes for patterning of the flower-like Asteraceae inflorescence. Plant Physiol. 2016;172:284–296. 10.1104/pp.16.00779

Zhong R, Lee C, Zhou J, McCarthy RL, Ye ZH. A battery of transcription factors involved in the regulation of secondary cell wall biosynthesis in Arabidopsis. Plant Cell. 2008;20:2763–2782. 10.1105/tpc.108.061325

Zhou Y, Yin M, Abbas F, Sun Y, Gao T, Yan F, et al. Classification and association analysis of Gerbera (Gerbera hybrida) flower color traits. Front Plant Sci. 2022;12:779288. 10.3389/fpls.2021.779288

Zoulias N, Duttke SHC, Garcês H, Spencer V, Kim M. The role of auxin in the pattern formation of the Asteraceae flower head (capitulum). Plant Physiol. 2019;179:391–401. 10.1104/pp.18.01119

